## Supporting Information for "Developmental and Aging Changes in Brain Network Switching Dynamics Revealed by EEG Phase Synchronization"

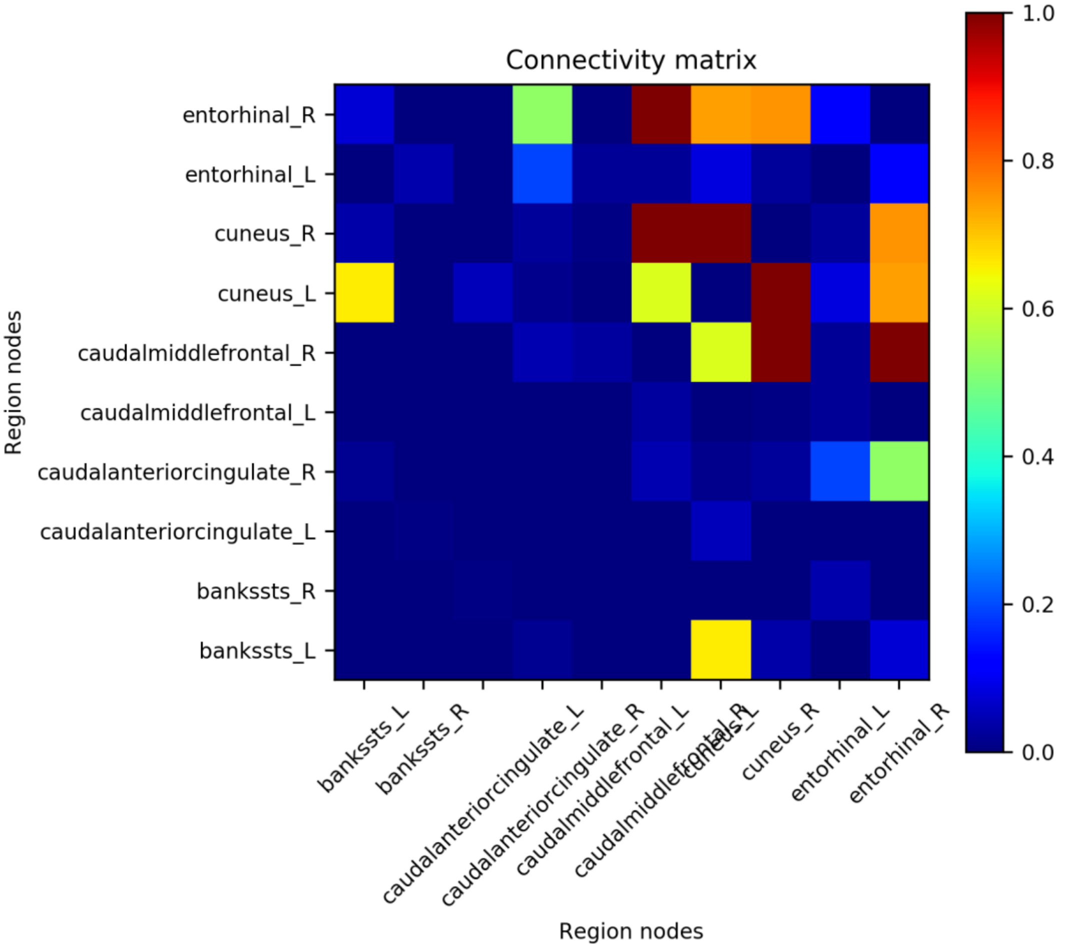


**Fig. S1. Connectivity used for simulations.**

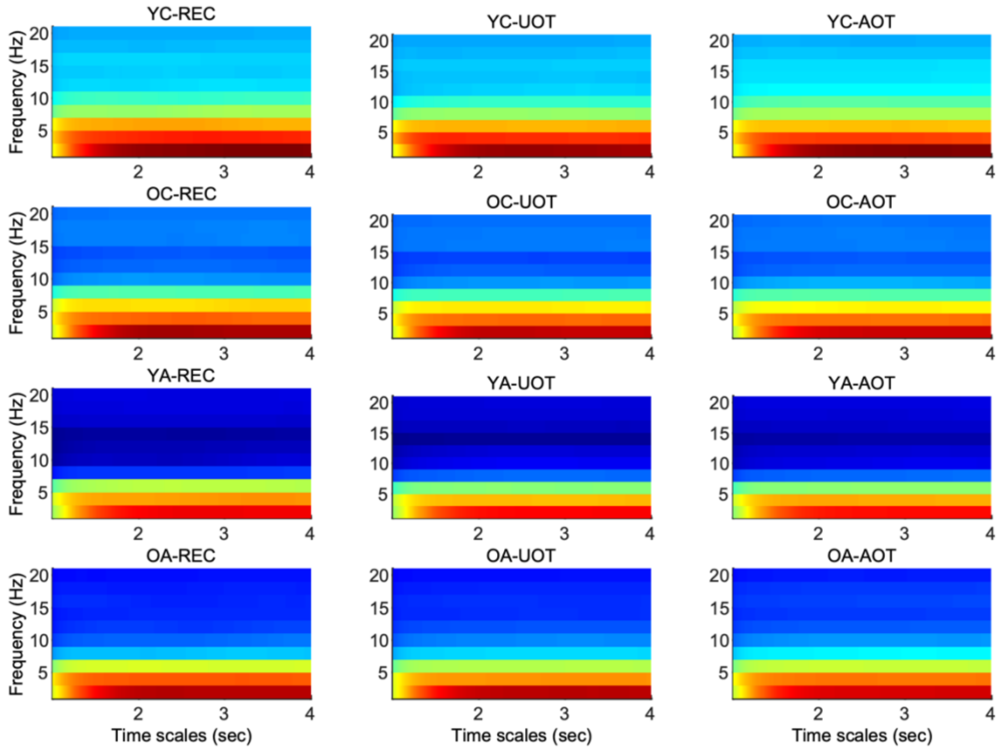


**Fig. S2. Group and condition means of $\mu_{JL(\tau,f)}$.** Groups are arranged in rows (age increases from top to bottom) and conditions in columns (attentional demands increase from left to right, i.e., '*REC*', '*UOT*', '*AOT*'). Time scale in seconds for all x-axes and frequency in Hz for y-axes. Note the transient effect for approximately the first second that corresponds to the length of the –fully overlapping- sliding window of phase synchronization computation. Only data for time scales in the range 1.004-2 sec, almost 1 sec after the length of the overlapping window, were used for further statistical analysis.

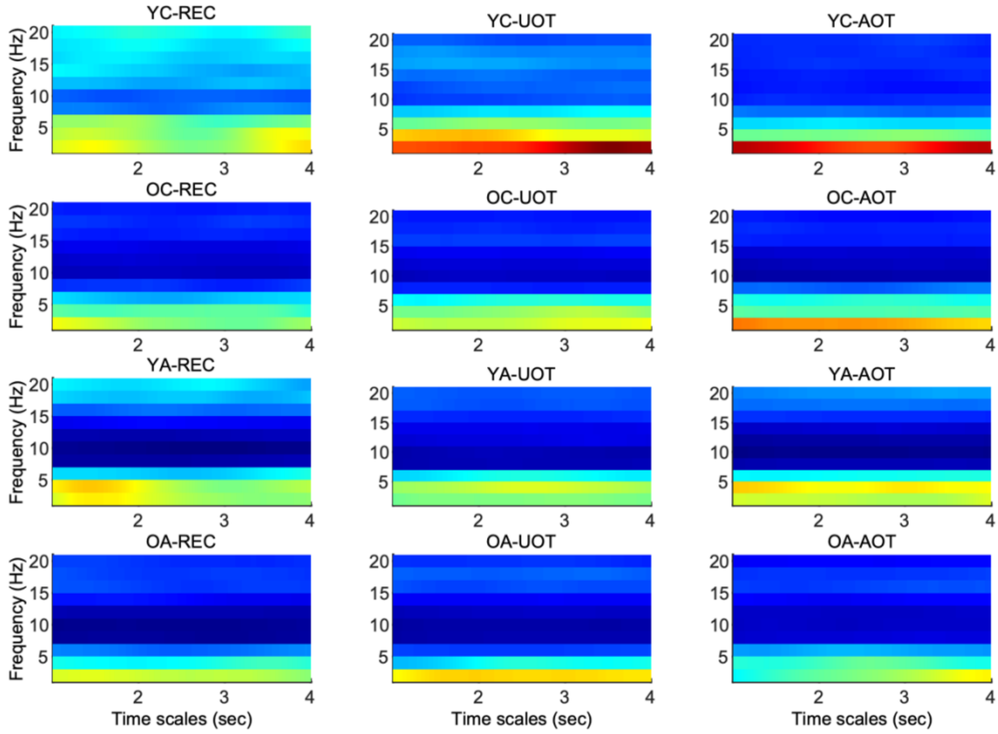


**Fig. S3. Group and condition t-score of $\mu_{JL(\tau,f)}$.** Figure arrangement and conventions similar to **Fig. S2**.

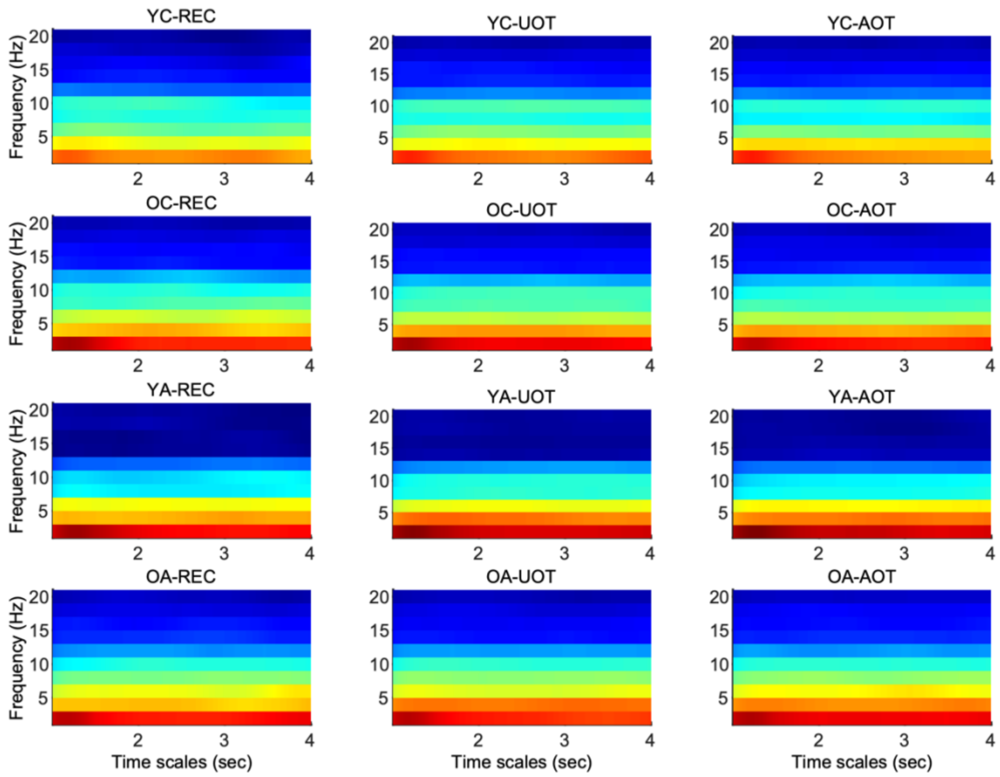


**Fig. S4. Group and condition means of $\sigma_{JL(\tau,f)}$.** Figure arrangement and conventions similar to **Fig. S2**.

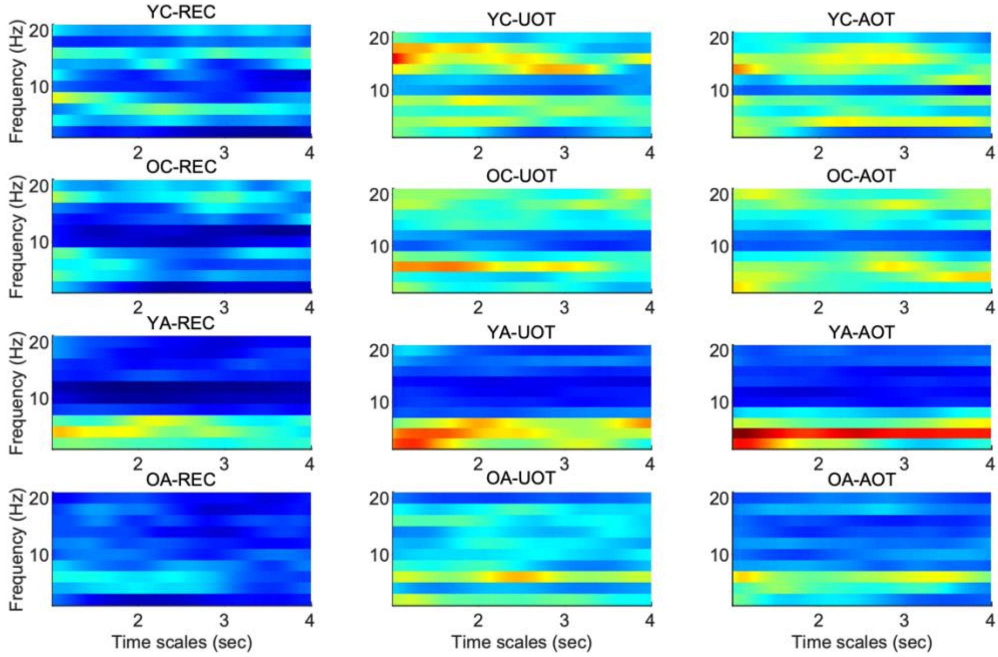


**Fig. S5. Group and condition t-score of $\sigma_{JL(\tau,f)}$.** Figure arrangement and conventions similar to **Fig. S2**.

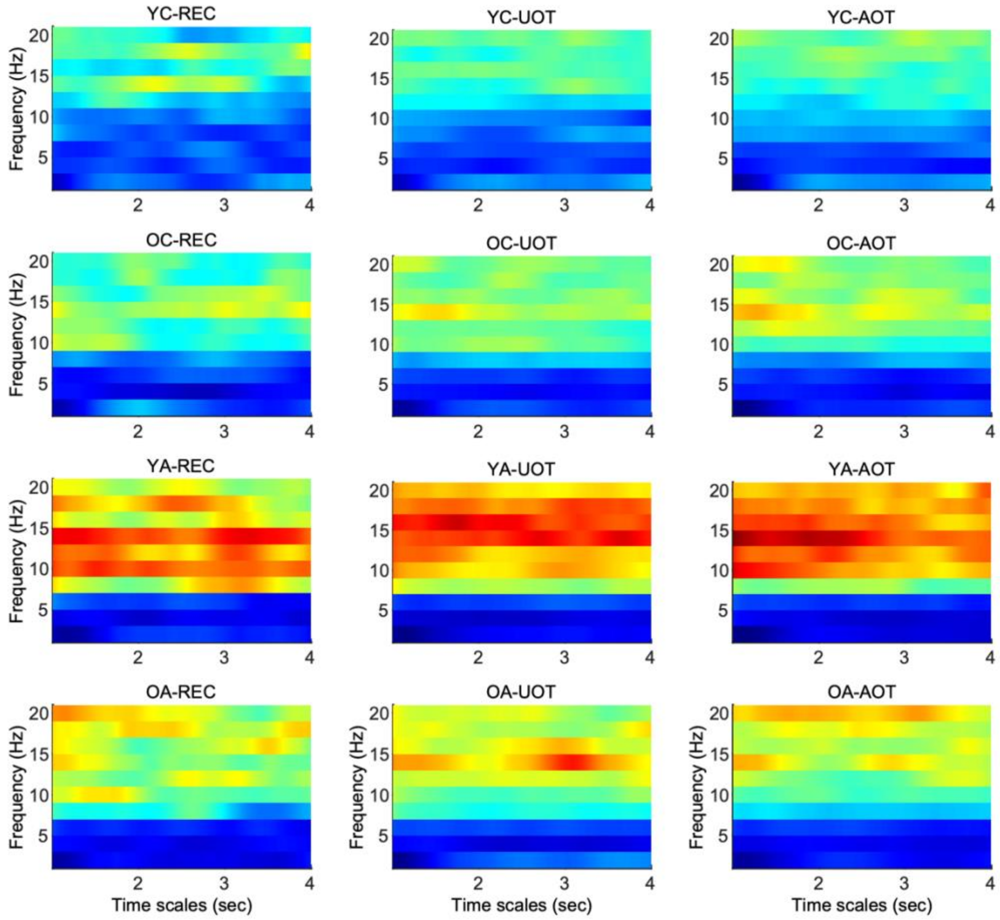


**Fig. S6. Group and condition means of $k_{JL(\tau,f)}$.** Figure arrangement and conventions similar to **Fig. S2**.

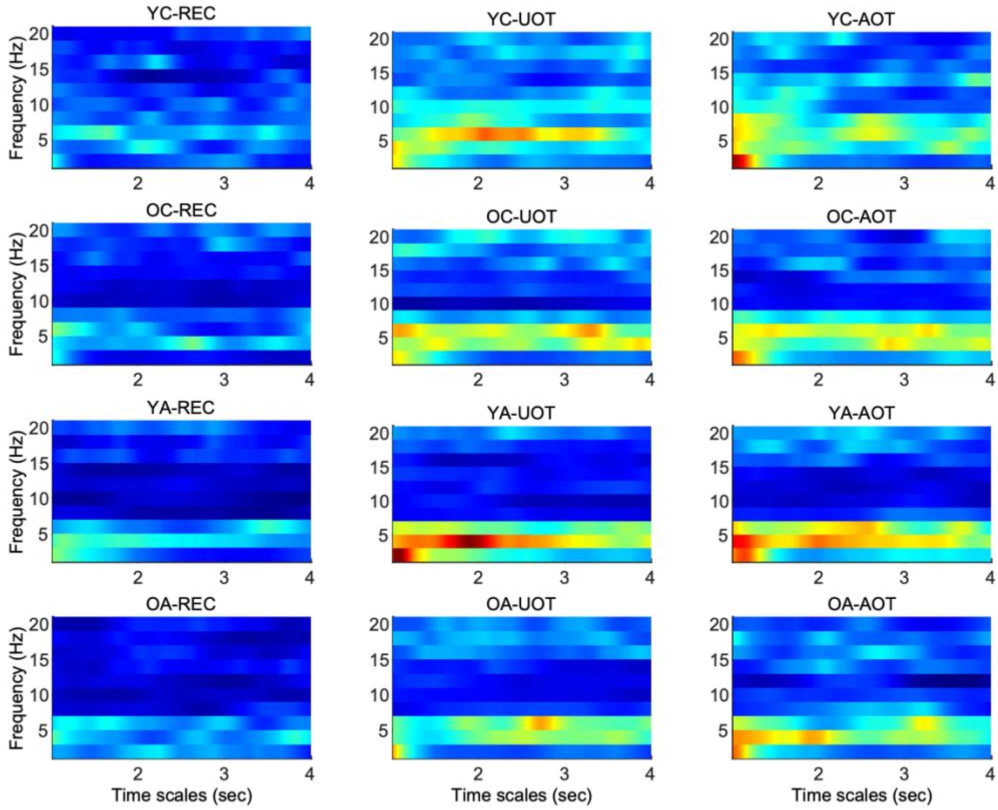


**Fig. S7. Group and condition t-score of $k_{JL(\tau,f)}$.** Figure arrangement and conventions similar to **Fig. S2**.

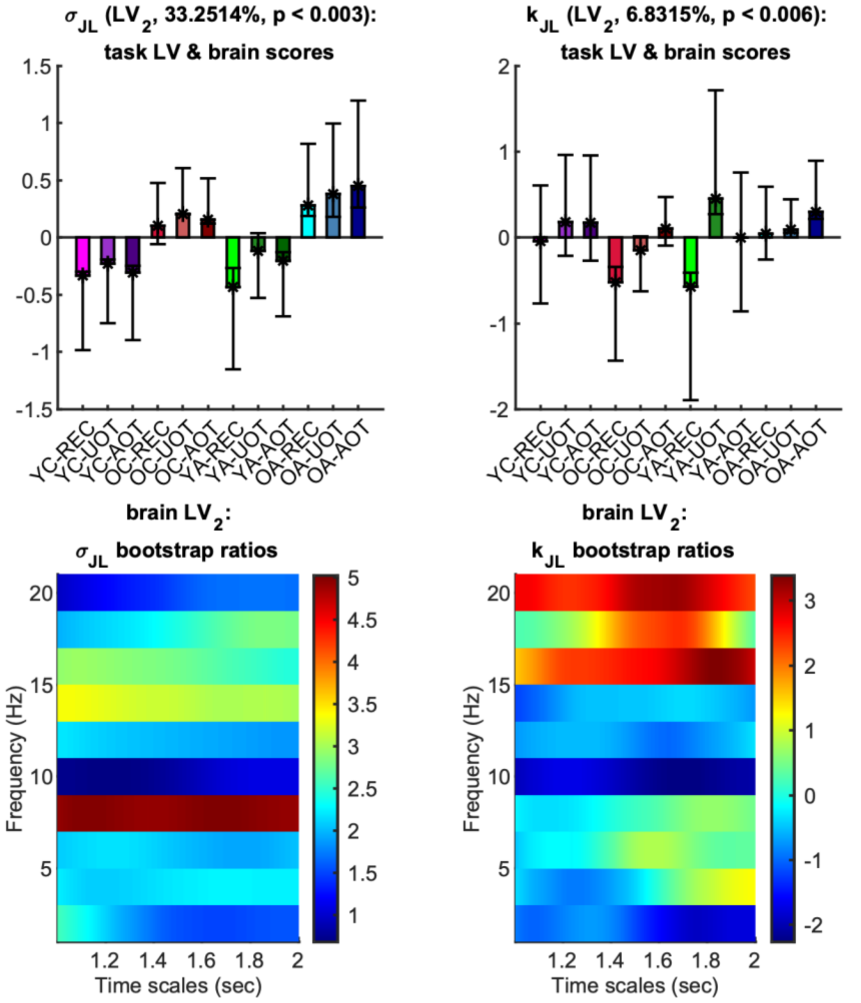


**Fig. S8. Second task latent variable, normalized brain scores and brain latent variable of the phase synchronization dynamics metrics.** *Top row:* Each panel shows the weights of the second *task latent* (asterisks), and the normalized brain scores with 95% confidence intervals (bars and errorbars) *variable* $\sigma_{JL(\tau,f)}$ (left) and $k_{JL(\tau,f)}$ (right). Color and other conventions are identical to *Figure 9* of the main text. This LV is significant ($p < 0.003$ for $\sigma_{JL(\tau,f)}$ and $p < 0.006$ for $k_{JL(\tau,f)}$) and explained approximately 33.3% and 6.8% of the covariance, respectively It contrasts '*OC*' and '*OA*' to '*YC*' and '*YA*' –in general- with statistical reliability (non-overlapping confidence intervals) for $\sigma_{JL(\tau,f)}$. Instead, the contrast described by the respective LV for $k_{JL(\tau,f)}$ is not reliable at all. *Bottom row:* Each panel shows the bootstrap ratios of the brain latent variable of $\sigma_{JL(\tau,f)}$ (left) and $k_{JL(\tau,f)}$ (right). Positive (negative) values correspond to elements that correlate positively (negatively) with contrasts of the top row, i.e., where that young adults (children) have higher (lower) values, respectively. Only values at 8 Hz for $\sigma_{JL(\tau,f)}$, and at 16 and 20 Hz for a short range of scales around 1.6 sec and 1.85 sec respectively, for $k_{JL(\tau,f)}$, have bootstrap ratios with absolute values greater than 2.5758. Therefore, these contrasts are statistically unreliable in general, and are not discussed in the main text.
